# MSMEG_1353 is a critical determinant of metabolic and cell envelope homeostasis in mycobacteria

**DOI:** 10.64898/2026.02.26.708182

**Authors:** Megha Sodani, Siddhesh Desai, Aaisha Ansari, Fiza Ahmed Shaikh, Nawab Singh Baghel, Pramod Kumar Gupta

## Abstract

The mycobacterial cell envelope is central to antibiotic resistance and stress adaptation, maintained by a network of essential proteins. Owing to their evolutionary divergence from other model organisms, mycobacteria encode numerous genus-specific genes that remain functionally uncharacterized, including the network of these essential genes. Here, we define MSMEG_1353 as a previously uncharacterized, essential ATP binding protein that plays a role in mycobacterial metabolism and cell envelope integrity. Conditional depletion of MSMEG_1353 resulted in severe growth attenuation, and the loss of characteristic mycobacterial cording. Further, the knockdown strain exhibited altered cell envelope permeability and an unusual antibiotic susceptibility profile, marked by isoniazid resistance accompanied with hypersensitivity to vancomycin. Functional analysis revealed pronounced membrane depolarisation and severe envelope defects, including distortion of outer membrane layers. Concomitantly, integrated transcriptomic and metabolomic profiling demonstrated widespread suppression of carbon metabolism accompanied by cofactor imbalance and redox perturbation, indicating a global metabolic rerouting upon MSMEG_1353 depletion. At the level of cell envelope composition, the lipid profiles indicate depletion of phthiocerol dimycocerosate (PDIM), trehalose polyphleate (TPP), and elevated levels of mycolic acid in the knockdown strain. Further, localisation studies revealed that MSMEG_1353 forms discrete clusters along the cell, with preferential enrichment at the old pole.

Collectively, our findings establish MSMEG_1353 as a critical determinant of metabolic homeostasis and envelope permeability in *Mycobacterium smegmatis*, linking central carbon metabolism, and adaptive cell envelope remodeling.

## 1. Introduction

Cell wall synthesis and carbon metabolism are closely linked pathways in a cell (1), both of which play critical roles in regulating bacterial growth. Their coordinated regulation is essential for maintaining growth, structural integrity, and survival. Mycobacteria possess a cell wall architecture and a mode of growth that are fundamentally distinct from those of well characterized model organisms and rely upon multiple specialized regulatory mechanisms (2). The mycobacterial cell envelope being one of the most structurally complex cell envelopes in bacteria, comprises a plasma membrane, a thick arabinogalactan-peptidoglycan complex, and an outer mycolic acid rich layer that forms the mycomembrane. This architecture renders mycobacteria intrinsically resistant to many antibiotics, restricts small molecule uptake, and supports environmental persistence (3–5). The asymmetric cell growth of bacteria is also one of the distinctive features of this organism whose implications and regulators remain to be studied (6, 7). Understanding the genetic determinants that regulate membrane homeostasis is therefore critical for deciphering mycobacterial physiology and identifying potential therapeutic targets. In the context of the escalating global burden of drug-resistant mycobacterial infections, elucidating the functions of essential genes has become imperative, not only to advance our understanding of pathogen physiology but also to enable the rational development of novel therapeutic interventions. Notably, three of the four primary first-line anti-tubercular drugs, target the mycobacterial cell wall. Advances in tuberculosis drug discovery further highlight cell envelope biosynthesis and metabolic pathways as highly tractable yet underexplored reservoirs for the development of novel anti-tubercular agents (8). Further, given the lipid rich nature of the mycobacterial cell envelope, defects in carbon metabolism are expected to impact the accumulation of lipid precursors, thereby influencing membrane architecture and cell wall integrity (9). Hence, a comprehensive dissection of the intricate architecture of the mycobacterial cell wall and its associated biosynthetic networks is therefore poised to reveal previously unappreciated vulnerabilities that may be exploited for drug discovery.

Fast-growing *Mycobacterium smegmatis* (*M. smegmatis*) is an established model for studying cell envelope biology (10). Utilizing this model organism, here we investigated MSMEG_1353, annotated as a hypothetical protein lacking experimental characterisation. MSMEG_1353 is essential for cell viability (11), which makes it difficult to study using conventional genetic approaches. However, the advent of CRISPRi technology has opened new avenues for the functional analysis of essential genes in bacteria (12), thereby accelerating research toward the identification of potential therapeutic drug targets (13, 14). While the functional significance of this gene remains unclear, a prior study has identified FtsQ as a binding partner of MSMEG_1353 (15), providing an initial clue toward its involvement in cell growth and division linked pathways. Given the close interconnection between cell division and cell wall biosynthesis, this observation prompted us to further investigate its functional role. Here, we combined CRISPRi-mediated knockdown with physiological, biochemical,-omics and microscopy based assays to examine how depletion of MSMEG_1353 affects cell envelope structure, permeability, and metabolism. Our results reveal that MSMEG_1353 plays an essential role in maintaining proper membrane architecture and metabolic homeostasis in the cell.

## 2. Results

### 2.1 MSMEG_1353 is a conserved ATP binding protein positioned at the interface of mycolic acid associated biosynthetic genes

To investigate the functional role of MSMEG_1353, an *in-silico* approach was employed to gain initial insights into its potential function. Bioinformatics analysis revealed that this gene is highly conserved across both slow and fast-growing mycobacterial species (Fig. 1A), including the obligate intracellular pathogen *Mycobacterium leprae* (*M. leprae*) which possesses a highly reduced genome with numerous inactivated pseudogenes that are no longer required for *in vivo* survival (16, 17). In *M. smegmatis*, MSMEG_1353 is situated within a genomic neighborhood composed largely of uncharacterized genes, and this local gene organisation does not exhibit a close synteny with the corresponding regions in *Mycobacterium tuberculosis* (*M. tuberculosis*) or *M. leprae*, except for a close by adjacent neighboring gene, MSMEG_1352 (LipG) which shares synteny across other species and has been characterized as a thioesterase/phospholipase primarily involved in phospholipid metabolism (18). Notably, however, MSMEG_1353 has two neighboring genes, MSMEG_1350 and MSMEG_1351 which retain homology to a well-characterized mycolic acid cyclopropane synthase, providing a conserved link to mycolic acid associated processes. Interestingly, the homologous locus in *M. tuberculosis* (Rv0647c) and *M. leprae* (Ml1898) is embedded within a genomic region enriched for genes involved in mycolic acid modification including *mmaA* (Rv0642-0645c), and *fabD2* (Rv0649) which is a malonyl-CoA:ACP transacylase that generates malonyl-ACP, a key precursor for fatty acid and complex lipid biosynthesis, especially important in mycobacterial cell wall lipid pathways (19). Thus, although the immediate neighboring genes differ across species, they consistently encode proteins involved in mycolic acid and lipid associated processes. This conservation of functional theme, despite divergence in local gene composition, strongly suggests that the MSMEG_1353 homolog operates within a lipid and cell envelope centric network across mycobacteria. In totality, the genomic neighborhood strongly shows association of this gene in lipid metabolism and its conservation across mycobacterium species, particularly *M. leprae* points towards an important role of this gene in mycobacterium growth and physiology.

**Fig. 1:**
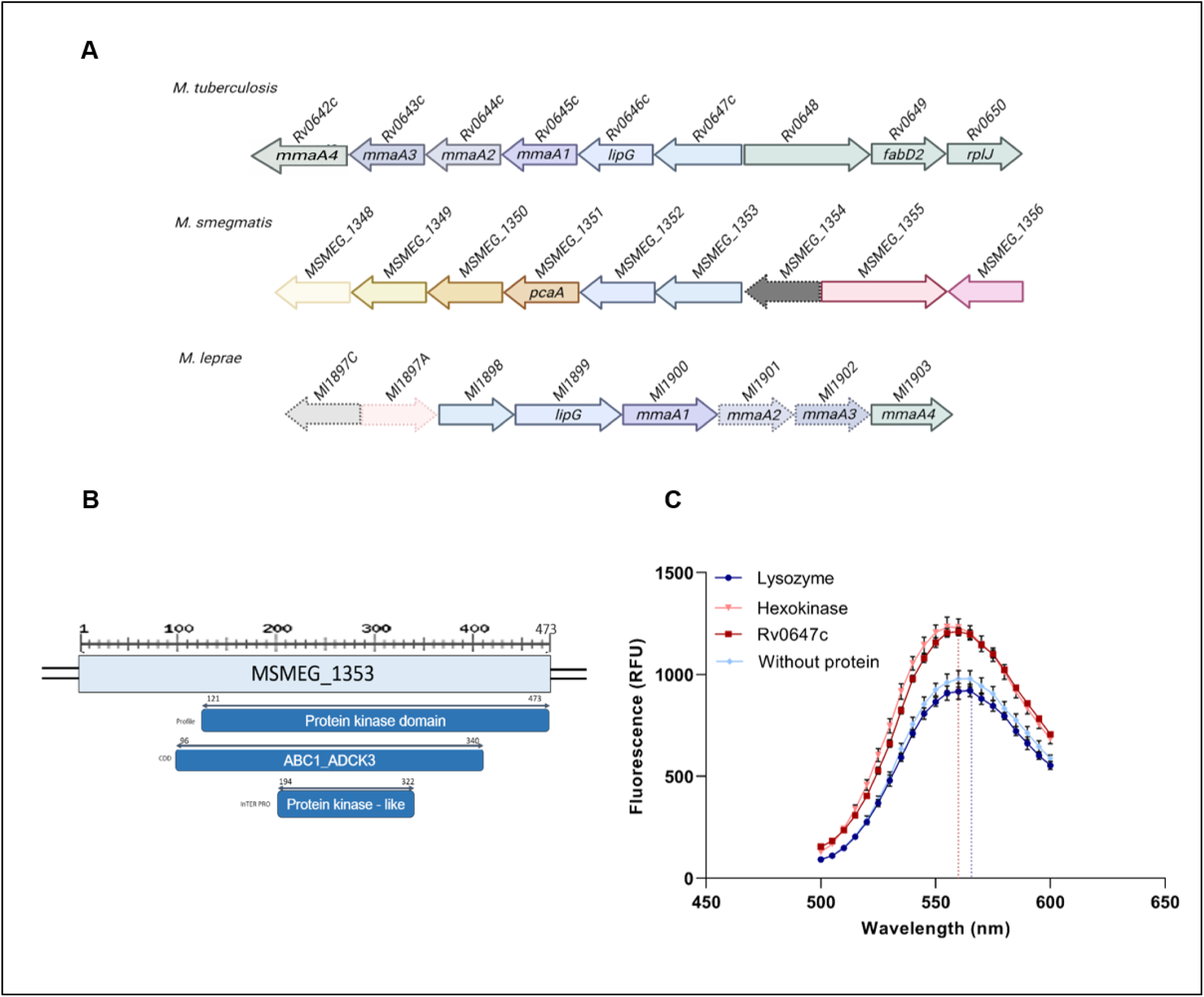
MSMEG_1353 encodes a conserved ATP-binding protein in mycobacteria. **A.** Genomic organisation of MSMEG_1353 and its homologues across different *Mycobacterium* species. (Arrow orientation indicates the direction of transcription, and boxed genes denote annotated pseudogenes). **B.** Schematic representation of the domain architecture of MSMEG_1353, highlighting the presence of an ATP-binding domain. **C.** Spectral scan of the TNP-ATP binding assay demonstrating ATP binding by MSMEG_1353 homolog, Rv0647c. The blue shift is elucidated as dotted lines.

Further, protein domain prediction using Pfam/InterPro algorithm revealed the presence of a conserved ATP binding domain (Fig. 1B) (20). Although the domain lacks typical homology to canonical kinases, it shares significant hits with ABC1 like atypical kinases. MSMEG_1353 shares ∼85% sequence identity with its *M. tuberculosis* homolog Rv0647c (S.1A), indicating high sequence conservation (21, 22). Leveraging the strong sequence homology and commercial availability of recombinant Rv0647c (CSB-EP308750MVZ), the predicted ligand interaction was validated with the TNP-ATP (2’,3’-O-(2,4,6-trinitrophenyl)adenosine-5’-triphosphate) fluorescence ATP binding assay (Fig. 1C). Here, TNP acts as a fluorophore and free TNP-ATP exhibits very low fluorescence in solution, whereas binding to a protein results in a marked increase in fluorescence accompanied by a characteristic blue shift (23). The assay included hexokinase, a bona fide ATP-binding protein, as a positive control, while lysozyme, which is not known to bind ATP, served as a negative control. Upon addition of TNP-ATP to Rv0647c, a clear enhancement in fluorescence along with a modest ∼5 nm blue shift was observed, confirming direct ATP binding activity (Fig. 1C). Collectively, these results indicate that MSMEG_1353 likely functions as an ATP interacting protein, involved in energy dependent processes in the cell.

### 2.2 CRISPRi-mediated knockdown of MSMEG_1353 causes growth attenuation and marked changes in colony features and ultrastructural defects in the cell envelope

To probe the physiological role of MSMEG_1353, a CRISPRi knockdown (KD) strain (24) and a pMV261-GFP fused-MSMEG_1353 over-expression strain (OE) was employed. A control strain, termed NTA, was used for comparison; this strain harbors the Cas12 effector gene but lacks a guide RNA targeting the mycobacterial genome. The KD strain was regulated by an inducible Tet promoter with anhydrotetracycline (ATc) serving as the inducer. Expression of MSMEG_1353 in the engineered strains were validated with the qPCR results, which confirmed the depletion of ∼167 fold and over-expression of ∼27 fold of MSMEG_1353 in the KD and OE strain, respectively (Fig. 2A). Upon induction, the KD strain displayed a reproducible and significant growth defect compared to the control NTA strain, most evident during exponential phase (Fig. 2B). The KD strain entered log phase later than the control strain or over-expressed strains but managed to reach comparable maximal OD values in ∼40 hours. This phenotype suggests that MSMEG_1353 contributes to optimal growth, particularly during rapid metabolic activity. On the other hand, the over-expression strain, 1353 OE continued to grow at par with the control pMV261-GFP strain (Fig. 2B).

**Fig. 2:**
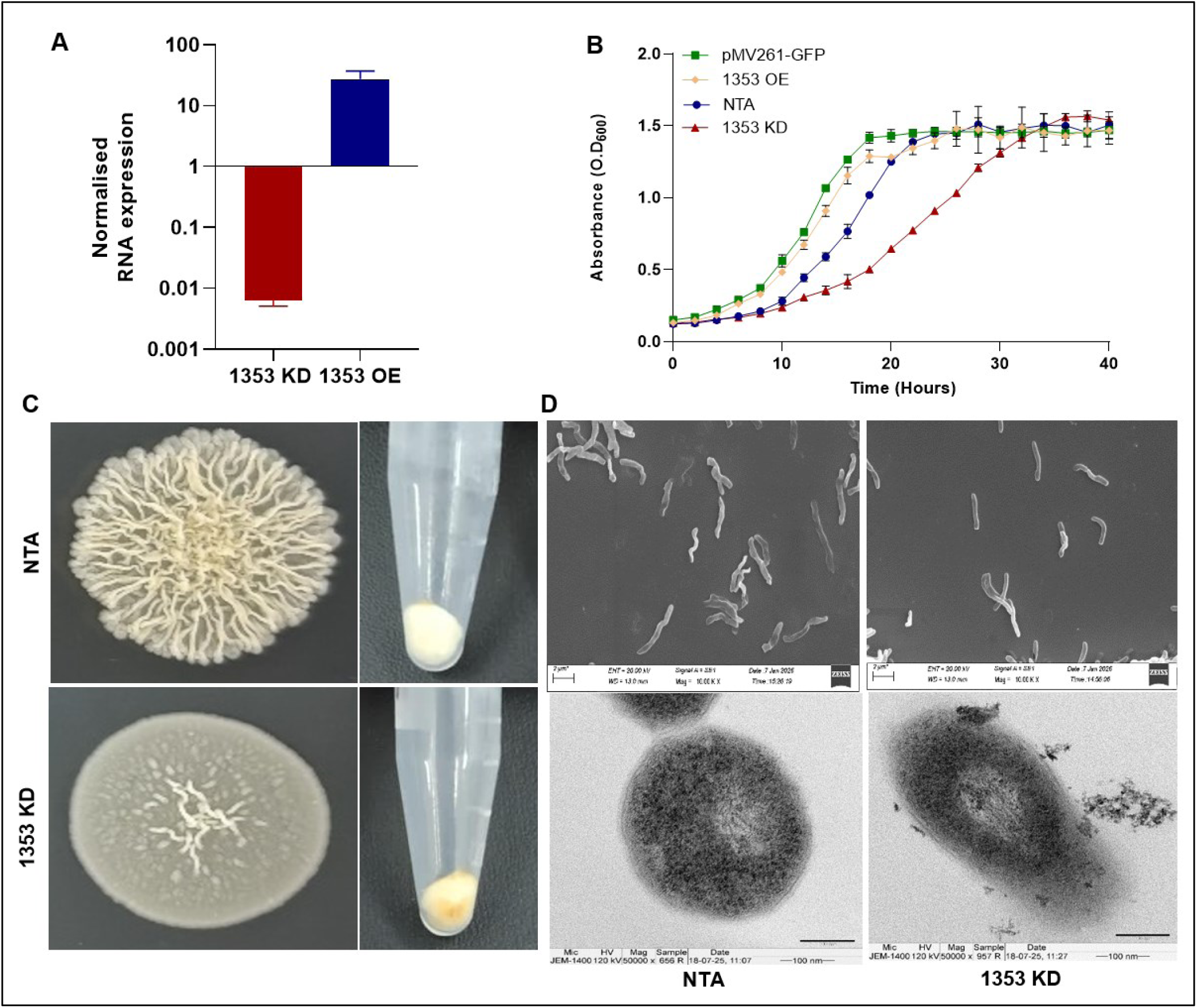
Knockdown of MSMEG_1353 alters gene expression, growth, colony and membrane characteristics. **A.** Relative quantification of MSMEG_1353 mRNA levels in the indicated strains, as determined by quantitative RT-PCR. **B.** Comparative growth kinetics of the indicated strains under standard culture conditions. **C.** Representative images showing differences in colony morphology, and cell pellet coloration between the MSMEG_1353 knockdown (KD) strain and the control NTA strain, as highlighted. **D.** Scanning electron micrographs (top panel) highlighting the morphological features of the control NTA strain and the MSMEG_1353 knockdown strain. Transmission electron micrographs (bottom panel) showing alterations in the cell envelope of the MSMEG_1353 knockdown strain compared to the control NTA strain.

Depletion of MSMEG_1353 led to pronounced changes in colony morphology (Fig. 2C). As expected, the NTA control colonies were rough with serrated edges typical of *M. smegmatis*, however, the KD strain produced smoother, round colonies with non-serrated margins. These morphological changes are often associated with alterations in the outer cell envelope (25, 26). In addition, cell pellets from the KD strain often exhibited a yellowish-orange coloration, whereas the control appeared cream/off-white (Fig. 2C), indicative of elevated carotenoid levels within the KD strain (27). Increased carotenoid production is known to impart the yellowish coloration and plays a protective role in quenching reactive oxygen species (ROS) and stabilizing lipid membranes, suggesting the presence of underlying redox imbalance or membrane stress in the KD strain (28).

The impact of MSMEG_1353 depletion on single cell level was examined using electron microscopy. The KD and the control strains were visualized under SEM and TEM to assess the surface and membrane architecture, respectively. In the SEM micrographs, the KD strain looked similar to control strain with no morphological defect as no detectable changes were observed in cell shape, or length (Fig 2D, top panel), implying that the knockdown of MSMEG_1353 does not lead to any significant changes in the size or the shape of strain. Interestingly, in the TEM micrographs, control cells displayed the characteristic triple layered envelope with a well-defined, electron dense outer membrane. However, the KD cells showed dramatic membrane defects, including shedding and peeling of the outer membrane and irregular and distorted membrane layers (Fig. 2D, bottom panel). The membrane showed electron lucent regions indicating weakened structural integrity and areas suggestive of cytosolic leakage, consistent with loss of envelope containment. These abnormalities indicate that MSMEG_1353 is essential for maintaining proper organisation or stability of the mycomembrane.

### 2.3 Membrane permeability and membrane potential is significantly altered in MSMEG_1353 KD strain

The impact of repression of MSMEG_1353 on membrane permeability was further probed by SDS disc diffusion assay and the susceptibility to the surfactant was scored in terms of the zone of inhibition (ZOI). SDS is a well-known anionic surfactant, routinely used to test mycobacterium membrane integrity (29). Upon exposure to 5% SDS, MSMEG_1353 KD strain showed a significantly larger ZOI compared to control, NTA strain (Fig. 3A and S.1B). While MSMEG_1353 KD exhibited a ZOI of 32 ± 2 mm, the NTA ZOI was measured at 29 ± 1 mm (average of three independent readings). As observed, MSMEG_1353 KD strain was significantly more susceptible to SDS stress than the NTA strain indicating alterations in membrane permeability.

**Fig. 3:**
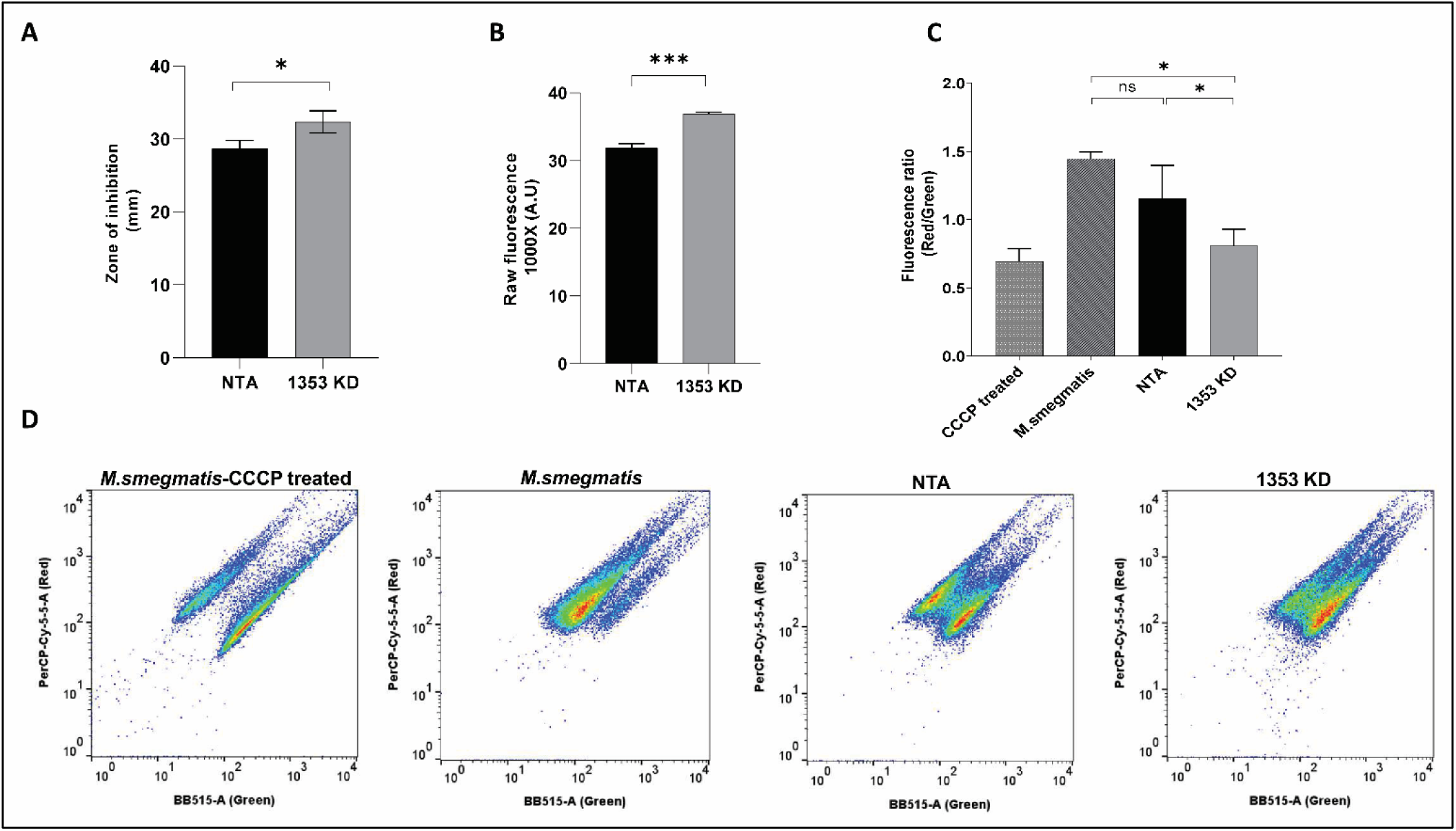
Knockdown of MSMEG_1353 alters membrane physiology. **A.** SDS disc assay bar graph showing zone of inhibition of MSMEG_1353 KD and NTA strain. **B.** Bar graph elucidating the ROS levels as evidenced by raw fluorescence levels of DCF observed in MSMEG_1353 KD and control NTA. **C.** Bar graph showing differential membrane potential, expressed as fluorescence ratio (red/green) of DiOC2(3), of intact and CCCP treated mycobacterium wild type strain, MSMEG_1353 KD and NTA strains. **D.** Dot blots showing the green and red population in correspondence with the membrane potential of the indicated strains.

Given the alterations observed in cell envelope permeability and integrity and their known influence in redox homeostasis (30), intracellular reactive oxygen species (ROS) levels were assessed using a standard DCFDA (2′,7′-Dichlorofluorescin diacetate) assay. Consistent with this, the KD strain exhibited significantly higher DCF (dichlorodihydrofluorescein) fluorescence compared to the control NTA, indicating elevated ROS levels upon MSMEG_1353 depletion (Fig. 3B). Taken together, these data suggest that depletion of MSMEG_1353 perturbs redox homeostasis and envelope stability in *M. smegmatis*.

Subsequently, the membrane physiology was investigated using a membrane potential sensitive dye, DiOC2(3) which was used to monitor the electric potential of the membrane in response to the depletion of the gene (31). DiOC2(3) exhibits green fluorescence in all bacterial cells, but the fluorescence shifts towards red emission as the dye molecules self-associate at the higher cytosolic concentrations caused by larger membrane potentials (32). As a positive control for the disruption of membrane potential (DiOC_2_(3) labeling), cells were incubated with carbonyl cyanide *m*-chlorophenyl hydrazone (CCCP), a strong uncoupler. The membrane potential was assessed by red to green fluorescence ratio, and it was found that the KD strain exhibited a marked reduction in membrane potential compared to control, consistent across biological replicates while the control strains NTA or wild type *M. smegmatis* showed no significant difference (Fig. 3C and D). Lower membrane potential is indicative of compromised proton motive force, often associated with membrane damage, ion leakage, or altered membrane protein function. This observation aligns with TEM data showing compromised membrane layers, reinforcing the notion that MSMEG_1353 is required for maintaining a functional and energized membrane.

### 2.4 Depletion of MSMEG_1353 results in selective alterations in antibiotic susceptibility

To investigate the influence of MSMEG_1353 on drug susceptibility, an antibiotic susceptibility profiling was carried out for the strains. Strikingly, the KD strain showed a severe resistance to isoniazid, an inhibitor of mycolic acid synthesis (Fig. 4A). Even at 10-fold higher concentration of MIC, where more than 90% control cells lose viability, MSMEG_1353 KD strain shows only a ∼50% susceptibility. The pronounced resistance to isoniazid indicates that the knockdown either impedes isoniazid access to its target or activates compensatory mechanisms that diminish drug efficacy. The latter explanation is more plausible, as isoniazid activation is NAD-dependent, and perturbations in cellular redox balance which are often observed during cell wall metabolic stress are likely to influence susceptibility to the antibiotic (33). Conversely, the KD strain displayed hypersensitivity to vancomycin (Fig. 4B), a large glycopeptide that cannot easily penetrate the intact mycomembrane of mycobacteria (34). Hence, increased vancomycin susceptibility in knockdown strain indicates heightened cell wall permeability. No significant changes were observed in susceptibility to ethambutol (Fig. 4C) or bedaquiline (Fig. 4D), indicating that the defect induced by MSMEG_1353 depletion affects select aspects of the envelope rather than general drug permeability. Together, these results demonstrate that MSMEG_1353 contributes towards maintaining a functional barrier that selectively modulates antibiotic access or efficacy.

**Fig. 4:**
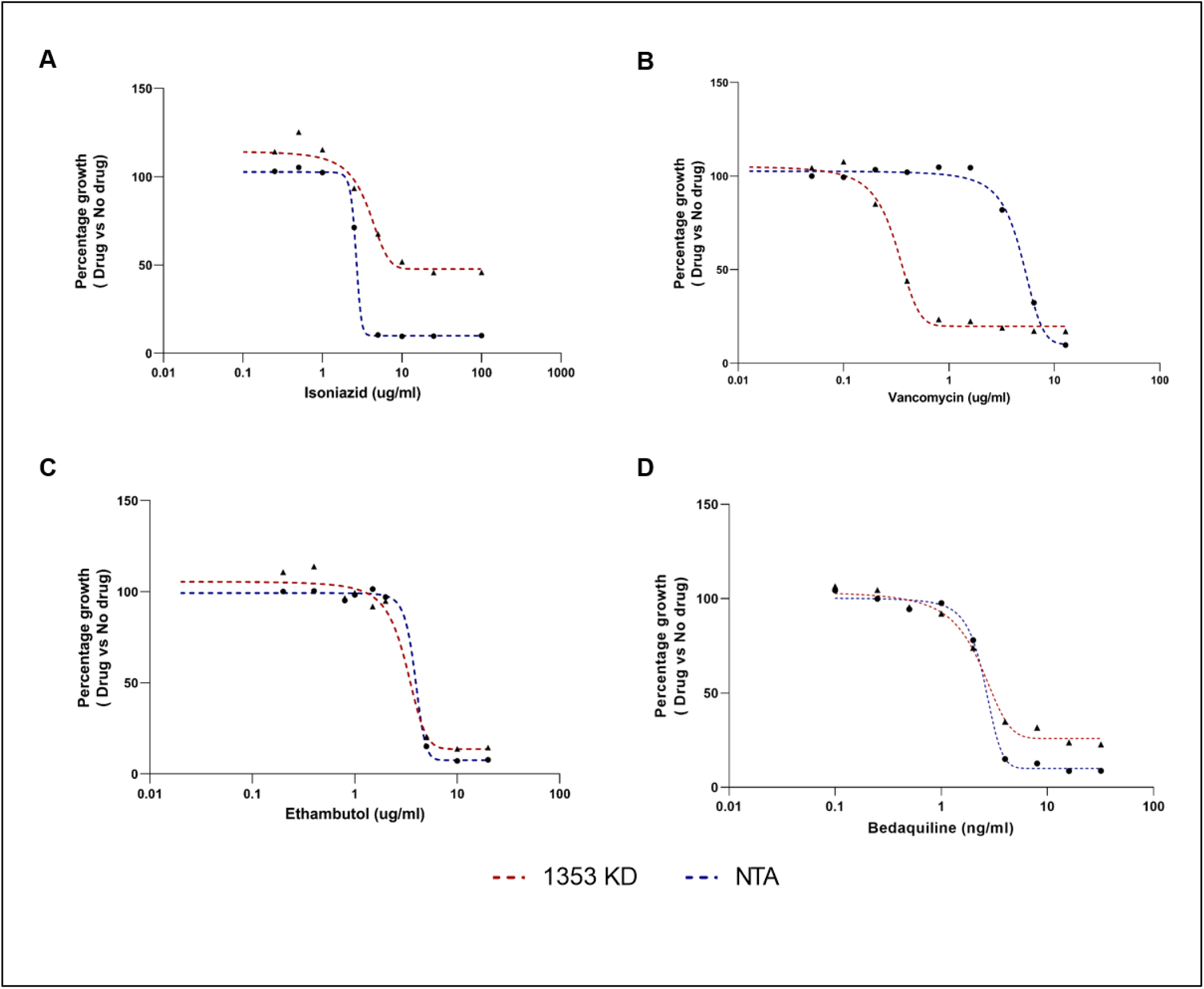
Antibiotic susceptibility of the MSMEG_1353 knockdown strain. A-D. Growth inhibition at varying concentrations of each antibiotic was assessed in broth culture. To account for the effect of gene repression itself, growth was normalized by expressing the percent growth in drug-containing media relative to growth in drug-free media at the corresponding concentrations.

### 2.5 MSMEG_1353 is a polar localised protein that accumulates preferentially at the old pole

To examine the localisation of MSMEG_1353, a C-terminal GFP-and FLAG-tagged fusion protein was expressed under a constitutive promoter, Phsp. Immuno-blot analysis using an anti-FLAG antibody confirmed a band at ∼80 kDa, corresponding to the expected size of the GFP-MSMEG_1353 fusion (S.1C). Fluorescence microscopy revealed that MSMEG_1353 does not distribute uniformly across the membrane unlike the GFP only strain (Fig. 5A). Instead, it forms discrete clusters along the cell, with pronounced enrichment at one of the two poles (Fig. 5A and 5B). Since mycobacteria grows asymmetrically, where one pole grows more than the other (35), a D-Amino acid analogue incorporation can be traced to identify the faster-(old) or slower-(new) growing pole. Hence, to determine pole identity, a RADA (D-amino acid analogue) pulse chase assay was set up where cells were uniformly grown in RADA at first and subsequently washed to chase the poles. Under these conditions, the slower growing new pole would now retain more RADA than the faster growing old pole. With pole identity thus established, the confocal fluorescent images showed that the GFP-MSMEG_1353 fluorescence intensity was significantly higher at the old pole, suggesting that MSMEG_1353 may function at faster growing regions of the cell (Fig. 5C). Polar localisation is characteristic of several envelope proteins involved in cell wall maturation, and membrane remodeling, further supporting the involvement of the gene towards growth and cell wall metabolism (36, 37).

**Fig. 5:**
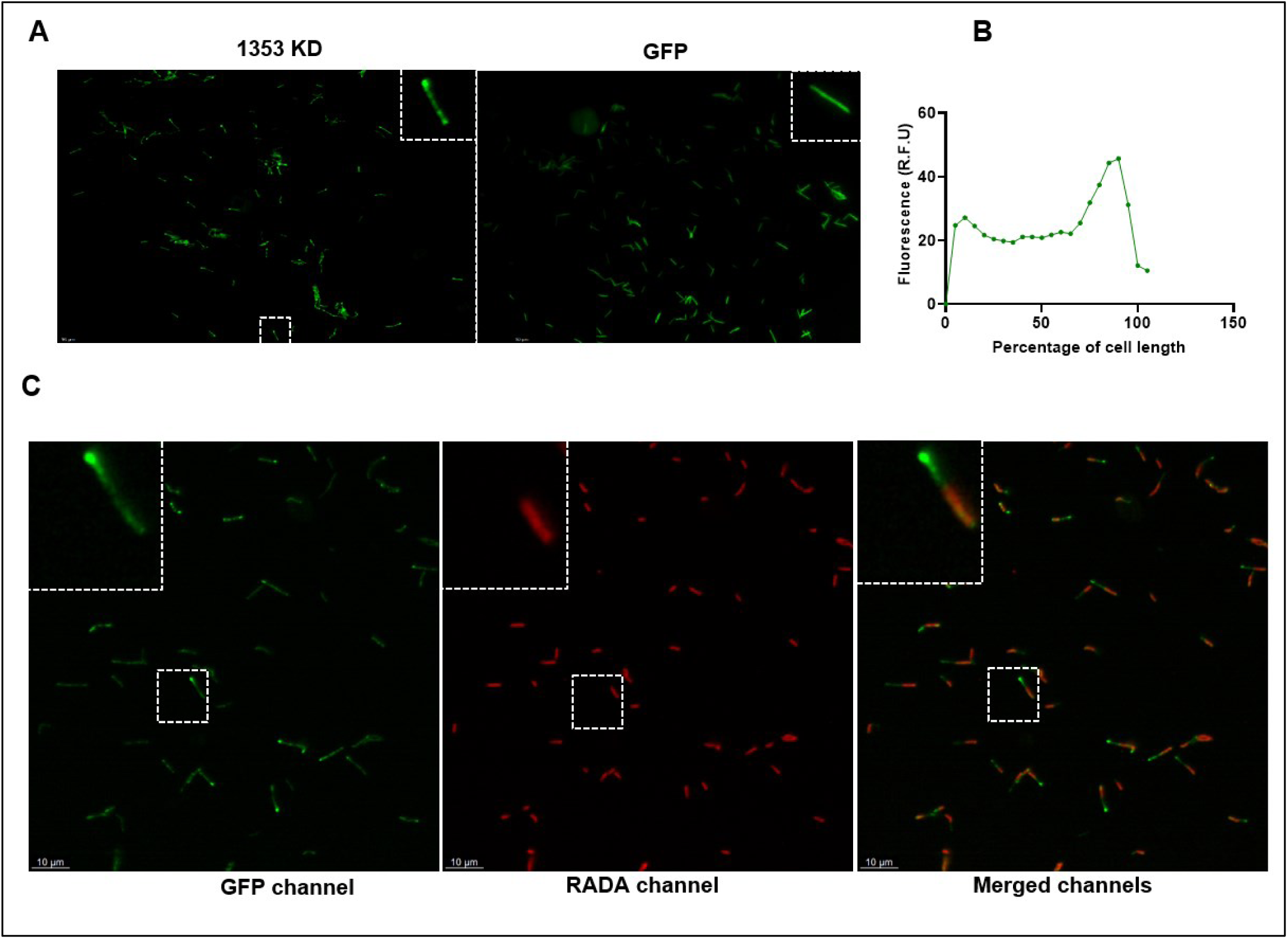
MSMEG_1353 localizes preferentially to the old cell pole. **A.** Fluorescence micrographs showing polar accumulation of MSMEG_1353-GFP, compared to uniform fluorescence in the GFP-only control. **B.** Fluorescence intensity profile of the MSMEG_1353-GFP strain along the length of the cell, indicating differential protein accumulation at the two poles (n=20). **C.** Fluorescence micrographs showing localisation of MSMEG_1353 at the growing old pole. left: GFP channel; middle: TRITC channel; right: merged image. The dotted box indicates the zoomed region displayed in the solid-lined boxes in the left corner.

### 2.6 Knockdown of MSMEG_1353 leads to global transcriptomic changes in the cell

Global RNA-sequencing analysis revealed extensive transcriptional reprogramming, marked by coordinated repression of growth associated metabolic pathways and concurrent induction of stress adaptive, transport, and regulatory functions. The significant genes were selected on the basis of P-adj value <= 0.05 and log2 fold change of +1 for up and-1 for down regulated. Based on this threshold, 156 genes were found to be up-regulated and 104 genes were observed to be down-regulated (Fig. 6A and S.2). The top 20 differentially expressed genes are highlighted in Fig. 6B. A few selected genes were verified with the qPCR data (Fig. 6C).

**Fig. 6:**
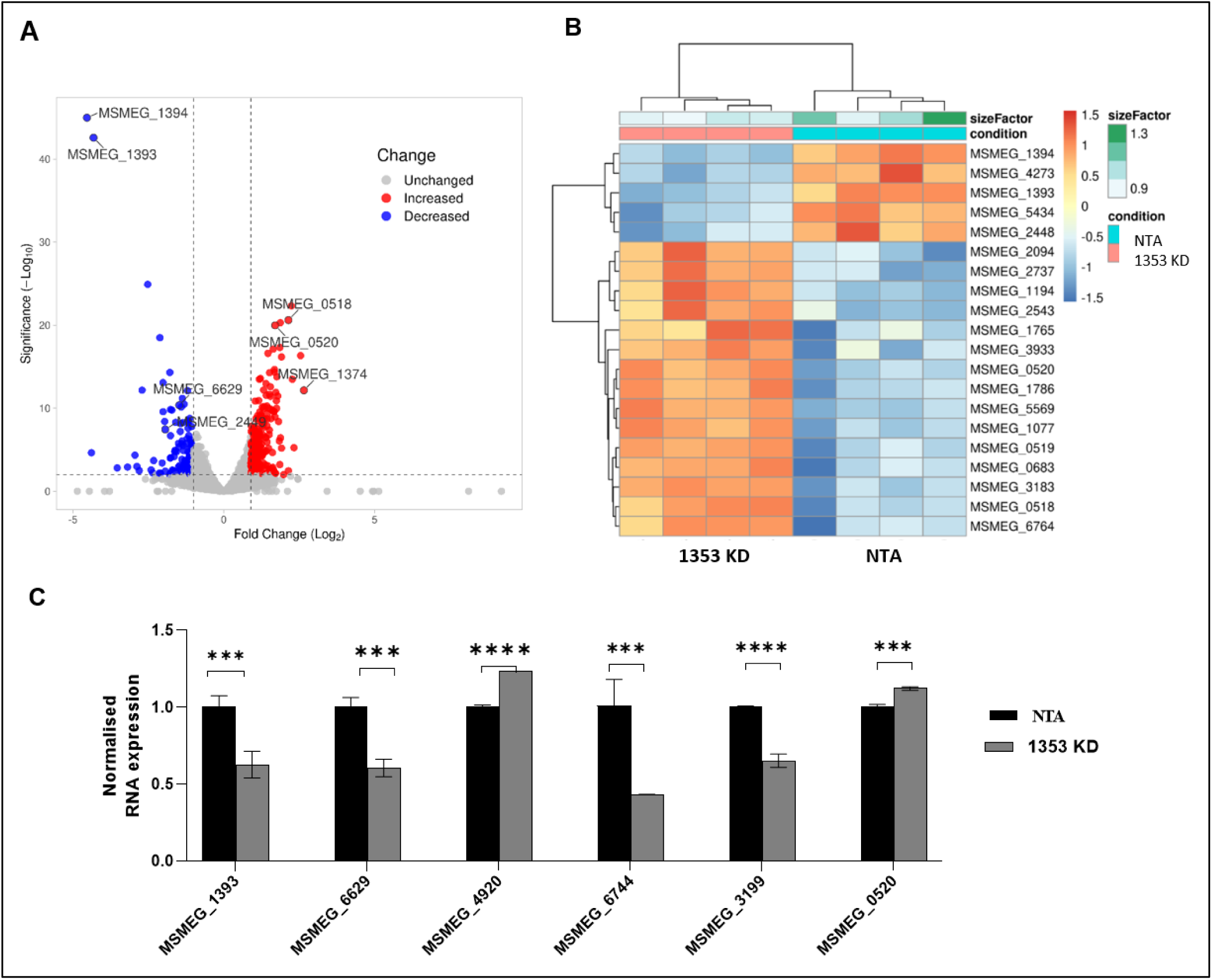
Transcriptome analysis of MSMEG_1353 KD strain. **A.** Volcano plot of differentially expressed genes where 104 genes with diminished expression and 156 genes with increased expression **B.** Heat map depicting expression of the top most differentially expressed genes in MSMEG_1353 KD strain vs NTA **C.** Quantitative analysis of RNA levels by qRT-PCR of some differentially expressed genes detected in RNASeq.

A prominent feature of the transcriptional landscape was the widespread down-regulation of central carbon metabolism. Key enzymes governing glycolysis and gluconeogenic flux, including pyruvate kinase (MSMEG_0154) and phosphoenolpyruvate synthase (MSMEG_3934), were significantly repressed, consistent with energy conservation under carbon limitation and stress. Notably, the most significantly down-regulated transcript in our RNA-Seq dataset was MSMEG_1394 (Fig. 6A and 6B), a gene that remains functionally uncharacterized in *M. smegmatis*. However, its homolog in *M. tuberculosis* has previously been reported to be down-regulated in a phosphoglucomutase mutant, implicating a potential link to central carbon metabolism (38). Multiple enzymes involved in short-chain and alternative carbon metabolism, such as methylmalonate-semialdehyde dehydrogenase (MSMEG_2449), malonate decarboxylase (MSMEG_6629), enoyl-CoA hydratase (MSMEG_3567), and 4-hydroxybutyrate CoA transferase (MSMEG_6310), were also reduced, further indicating a global contraction of metabolic flexibility (S.2). Further amino acid biosynthetic genes (e.g., phosphoglycerate dehydrogenase (MSMEG_6115), chorismate mutase (MSMEG_5536), methylmalonate-semialdehyde dehydrogenase (MSMEG_2449), and branched chain amino acid transport (MSMEG_3247) were also found to be repressed. Transcriptional repression extended to pathways supporting biosynthetic commitment and cofactor production. For example, quinolinate synthetase A (MSMEG_3199) and biotin synthase (MSMEG_3194) were down-regulated, pointing to diminished *de novo* synthesis of NAD and biotin, essential cofactors for redox metabolism (39) and fatty acid biosynthesis (40). In line with this, several oxidoreductases, including FMN (MSMEG_1424) and FAD/NAD-dependent dehydrogenases (MSMEG_6744) and succinate semialdehyde dehydrogenase (MSMEG_2488), were reduced, suggesting altered redox enzyme deployment.

In contrast to the widespread suppression of growth associated metabolism, a strong induction of transport, scavenging, and stress response pathways was also observed (S.2). Multiple ABC transporters, including those involved in glycerol-3-phosphate (MSMEG_0518), ribose (MSMEG_1374), ectoine (MSMEG_3548), were significantly upregulated, along with porins (MSMEG_0520) and major facilitator superfamily transporters (MSMEG_3586), indicating enhanced nutrient acquisition from the extracellular environment (Fig. 6B). Notably, glucose-6-phosphate isomerase (MSMEG_5541) was upregulated alongside ABC glycerol-3-phosphate transport, indicating attempted revival of upper glycolysis using scavenged carbons, suggesting uncoupling of upper and lower glycolysis due to downstream pyruvate kinase repression.

Starvation induced DNA protecting protein (MSMEG_6467) and uracil DNA glycosylase (MSMEG_2399) were induced, pointing to enhanced DNA protection and repair rather than replication. Several genes involved in cell envelope lipid metabolism, including mycocerosic acid synthase, polyketide cyclase, Ppm1, and 3-oxoacyl-[acyl-carrier-protein] reductase, were significantly up-regulated. This pattern is consistent with modifications of the cell surface, though the precise functional consequences remain to be elucidated.

Together, the RNA-sequencing data reveal a transcriptional program characterized by suppression of central metabolism, biosynthetic capacity, accompanied by activation of transport, stress response, and cell envelope remodeling pathways. This coordinated transcriptional architecture reflects a transition from a growth oriented state to a stress adapted, physiological program.

### 2.7 Metabolomics changes observed in MSMEG_1353 depleted strain

To capture the cellular level metabolic consequences of the MSMEG_1353 depleted condition, an untargeted metabolomic profiling was employed. Untargeted metabolomic profiling revealed widespread changes in intracellular metabolite pools, indicating that loss of MSMEG_1353 leads to a pronounced shift in metabolic state under the tested condition (S.3). Several metabolites could not be confidently assigned to known KEGG pathways, likely due to incomplete pathway curation, limited annotations, or identification constraints; these metabolites were manually removed from the analysis. Multivariate analysis showed clear separation between control and experimental samples, confirming robust and condition specific metabolic reprogramming in MSMEG_1353 KD (S.1D).

One of the dominant features of the metabolomics landscape was the global depletion of core metabolic and energy related metabolites. Several nucleotide intermediates, including coenzyme A (CoA), cytidine diphosphate (CDP), cytidine 5′-monophosphate (CMP), deoxyadenosine diphosphate (dADP), and deoxyguanosine diphosphate (dGDP), were significantly reduced (Fig. 7A and 7B). Consistent with this trend, several carbohydrate and sugar phosphate related metabolites, including 1,6-di-O-phosphono-α-D-glucopyranose and galactosamine, were diminished, reflecting reduced availability of carbohydrate derived intermediates and sugar-phosphate pools.

**Fig. 7:**
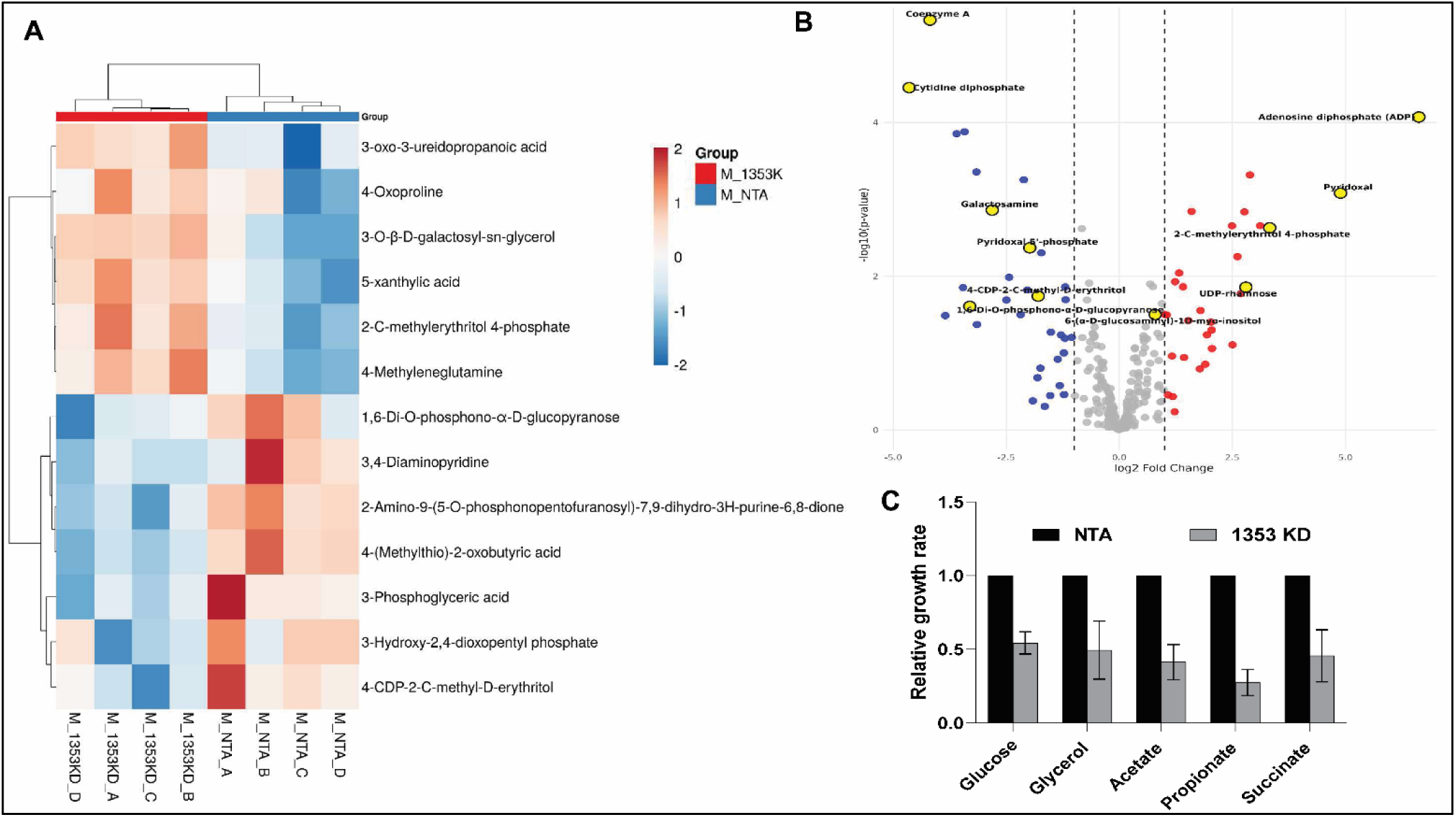
Metabolomics analysis of MSMEG_1353 KD strain. **A.** Heat map depicting expression of the top most differentially expressed genes across replicates in MSMEG_1353 KD strain vs NTA. **B.** Volcano plot of differentially expressed metabolites. **C.** Assesment of normalized specific growth rate of the indicated strains in presence of different carbon sources.

Metabolites associated with amino acid biosynthesis and nitrogen metabolism were broadly decreased. These included L-cystathionine, N-acetyl-L-citrulline, and the dipeptide L-alanyl-glutamine, indicating constrained amino acid pools and reduced anabolic capacity. Cofactor and vitamin associated metabolites were also markedly perturbed. Levels of the phosphorylated, cofactor-active forms of vitamin B6-pyridoxal 5′-phosphate and pyridoxine phosphate were reduced, while the non-phosphorylated precursors pyridoxal and pyridoxine accumulated, indicating disruption of vitamin B6 cofactor balance. The downstream non-mevalonate pathway intermediate 4-CDP-2-C-methyl-D-erythritol was also reduced, suggesting impaired downstream progression through isoprenoid-related biosynthetic steps.

In contrast to the widespread depletion of central metabolic intermediates, a distinct subset of metabolites accumulated, indicating selective enrichment of stress associated and turnover related processes. Energy and nucleotide recycling associated metabolites, including adenosine diphosphate (ADP), inorganic pyrophosphate, and ADP-ribose 1″,2″-cyclic phosphate, were elevated, consistent with altered cellular energy balance and enhanced nucleotide recycling. Several amino acid derived metabolites and dipeptides, including proline, N-acetylornithine, L-(+)citrulline, succinyl proline, glycyl-L-leucine, Ala-Glu dipeptide, and N(2)-succinyl-L-arginine, were also increased, reflecting enhanced protein turnover and nitrogen redistribution.

Metabolites associated with cell envelope remodeling and glycan related processes were enriched. Accumulation of UDP-rhamnose and 6-(α-D-glucosaminyl)-1D-myo-inositol indicates altered allocation towards cell wall-associated sugar nucleotide pools and glycosylated envelope components. Additionally, elevated levels of the upstream MEP pathway intermediate 2-C-methylerythritol-4-phosphate, together with depletion of the downstream CDP activated intermediate, suggest partial engagement of the pathway. This pattern is consistent with nucleotide and cofactor limitation.

To investigate the functional impact of the metabolic perturbations, we assessed the fitness of the KD strain relative to the wild type during growth on different carbon sources representing distinct entry points into central metabolism. The KD strain displayed the most pronounced fitness defect on propionate as the sole carbon source (Fig. 7C), exhibiting maximal growth attenuation and extended lag phase compared to the wild type, consistent with impaired detoxification of propionyl CoA or methylcitrate cycle dysfunction. An intermediate level of fitness loss was evident on glycerol, where relative specific growth rate was significantly reduced but less severely than on propionate (Fig. 7C). In contrast, growth on glucose, acetate, and succinate was comparably milder, with similar relative fitness impairments across these substrates that intersect glycolysis and the TCA cycle more directly. These carbon source specific vulnerabilities highlight a preferential reliance on robust anaplerotic flux and avoidance of propionyl CoA associated stress conditions in the KD strain, aligning with the CoA depletion pathway shifts observed in the metabolomics profile.

Collectively, MSMEG_1353 depletion induces global depletion of central intermediates, nucleotides, amino acids and cofactors, coupled with selective enrichment of envelope-remodeling metabolites. These perturbations align with compromised cell envelope integrity and supports MSMEG_1353’s role in supporting metabolomics homeostasis essential for robust growth and barrier function

### 2.8 Depletion of MSMEG_1353 leads to changes in cell envelope lipid components

To investigate the specific classes of cell envelope lipids affected by depletion of MSMEG_1353, we quantified major neutral and mycolate associated lipid species, including diacylglycerol (DAG), triacylglycerol (TAG), phthiocerol dimycocerosates (PDIM), trehalose monomycolate (TMM), trehalose dimycolate (TDM), trehalose polyphleates (TPP) and total mycolic acids. Mycobacteria produce a diverse repertoire of surface exposed lipids that play critical roles in cellular physiology and host interaction. Among these, TPPs are complex glycolipids associated with the mycobacterial cell surface (41). TPPs have previously been implicated in the formation of characteristic cording morphology in *Mycobacterium abscessus* (42). Consistent with this role, we observed a marked reduction in TPP levels in the MSMEG_1353 KD strain (Fig. 8A). This finding aligns with data from a genome wide CRISPRi chemical genomics screen, which identified MSMEG_1353 within a cluster of genes involved in TPP biosynthesis, further supporting a functional link of MSMEG_1353 influencing cell surface architecture (43). Depletion of MSMEG_1353 also resulted in a pronounced accumulation of mycolic acids (Fig. 8B). This increase in mycolic acid abundance in the knockdown strain is concordant with its enhanced resistance observed to isoniazid, a mycolic acid biosynthesis inhibitor. Further, PDIM is a critical, complex surface lipid found in the outer cell envelope of pathogenic mycobacterium and is known to compromise the cell wall permeability (44). In this context, PDIM levels were investigated and it was found to be selectively reduced in the KD strain, consistent with compromised outer envelope integrity and the observed hypersensitivity to vancomycin (45) (Fig. 8C). No significant changes were detected in DAG, TAG, TMM, or TDM levels across strains (Fig. 8D, 8E and 8F). By and large, the data indicate that MSMEG_1353 depletion elicits a broad perturbation of lipid homeostasis affecting the cell wall architecture in the cell.

**Fig. 8:**
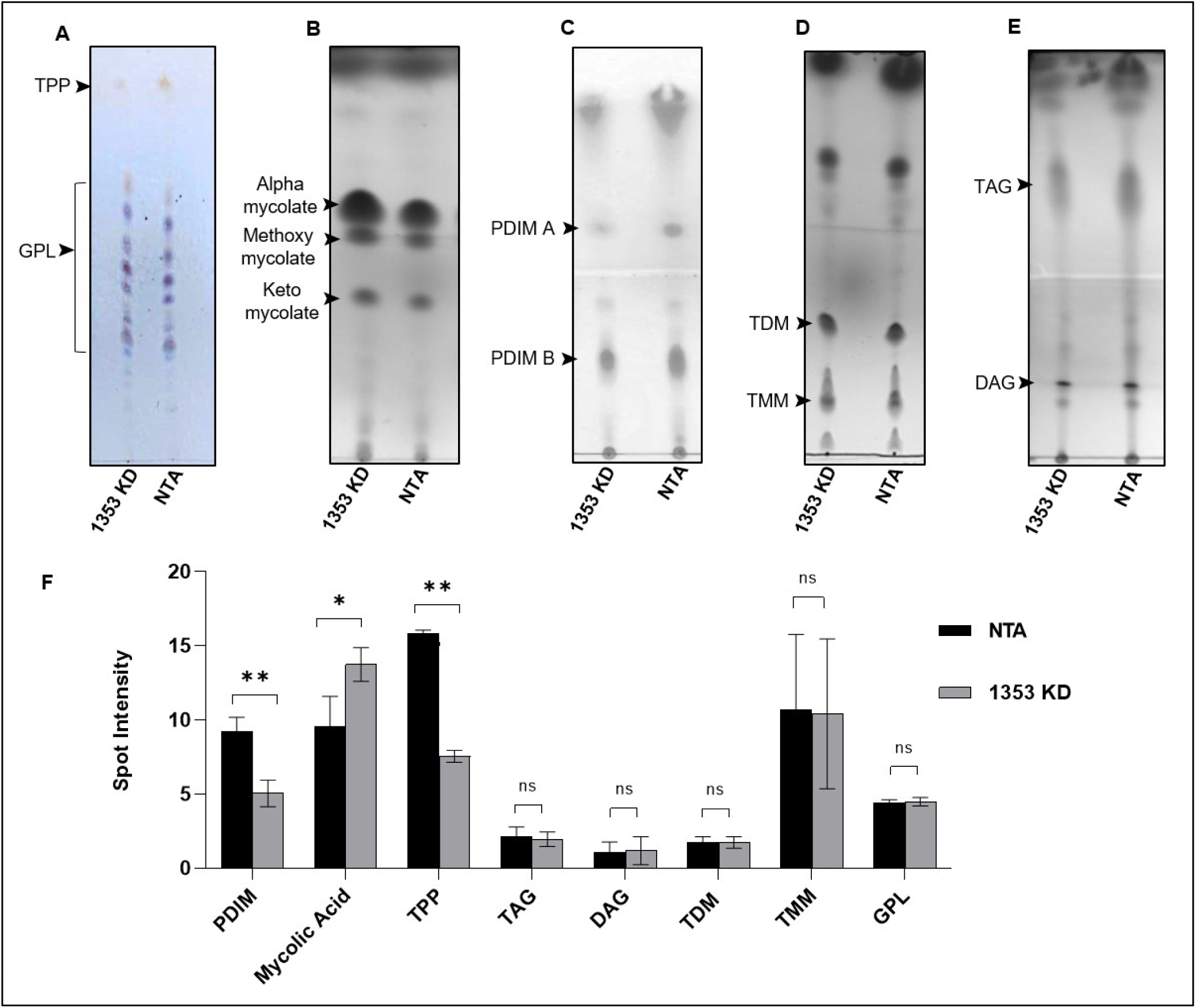
Depletion of MSMEG_1353 influences membrane lipid profiles in the cell. **A.** TLC analysis of trehalose polyphleates (TPP) and glycopeptidolipids (GPL) developed in chloroform:methanol (90:10, v/v) in MSMEG_1353 knockdown and NTA strains, visualized by anthrone staining in concentrated sulfuric acid (brown bands for TPP at higher Rf; blue bands for GPL). **B.** Thin-layer chromatography (TLC) analysis of mycolic acids in MSMEG_1353 knockdown and NTA strains, developed five times in hexane:ethyl acetate (19:1, v/v). Bands were visualized by iodine vapor exposure or charring. **C.** TLC analysis of apolar lipids (including PDIM) in MSMEG_1353 knockdown and NTA strains developed once in petroleum ether (40–60°C): diethyl ether (90:10, v/v). The major band corresponding to phthiocerol dimycocerosates (PDIM) is indicated. **D.** TLC analysis of trehalose mycolates (TMM/TDM) in MSMEG_1353 knockdown and NTA strains. Polar lipids enriched for trehalose monomycolate (TMM) and trehalose dimycolate (TDM) were developed in chloroform:methanol:water (90:10:1, v/v/v, visualized by phosphomolybdic acid staining by charring. **E.** TLC analysis of neutral lipids (TAG/DAG) in MSMEG_1353 knockdown and NTA strains developed in hexane:diethyl ether:acetic acid (70:30:1, v/v/v), visualized by phosphomolybdic acid staining or charring. **F.** Bar graph of densitometric analysis of all the TLCs depicted in A-E.

## 3. Discussion

Our understanding of the regulatory mechanisms governing cell envelope remodeling, and the intricate links between core metabolism and envelope integrity, remains at an early stage. However, recent studies have started to uncover functional connections between these processes (46–48). This gap in mechanistic understanding is particularly notable given that enzymes involved in cell envelope biosynthesis and remodeling constitute a rich reservoir of anti-bacterial drug targets. In this study, we investigate the function of the essential mycobacterial gene MSMEG_1353. The data collectively suggest that MSMEG_1353 likely functions as a metabolic-envelope coupling factor rather than serving as a classical biosynthetic enzyme affecting a specific class. The breadth of phenotypes arising from its depletion, spanning redox imbalance, membrane destabilisation, altered drug susceptibility, and transcriptional and metabolic reprogramming, argues against a single downstream defect and instead points to a failure of coordination between energy status and cell envelope maintenance.

A key conceptual insight emerging from this study is that the cell envelope integrity appears to be actively regulated by cellular metabolic state. This interpretation is consistent with recent reports demonstrating active regulation of mycomembrane integrity by cellular metabolic state, thereby supporting the notion that envelope architecture is tightly integrated with intracellular energetics (49–51). Mycobacteria invest a substantial fraction of their biosynthetic and energetic capacity into maintaining the lipid rich outer membrane, and it is reasonable to expect that any disruption in ATP availability, redox balance, or metabolite flux is likely to be sensed and compensated for. Indeed, in mycobacteria, such metabolic surveillance mechanism is frequently mediated by ATP binding proteins, including eukaryotic-like Ser/Thr protein kinases (STPKs), which function to couple cellular energy status with cell envelope biogenesis, growth, and stress responses (52–54). Within this framework, the ATP binding capability of MSMEG_1353, suggests that it may act as a molecular sensor or scaffold that links intracellular energy status to envelope associated processes. Rather than driving a specific reaction, MSMEG_1353 may stabilise or regulate multi-protein complexes involved in lipid handling, membrane assembly, or redox balancing at the membrane interface. Importantly, the polar localisation of MSMEG_1353 further supports this interpretation. Polar regions in mycobacteria are hubs of active cell envelope synthesis, where metabolic flux, and membrane assembly, converge. Localisation of MSMEG_1353 to the poles positions it ideally to integrate intracellular energy status with envelope associated processes, potentially by stabilizing membrane assembly at sites of active growth.

The envelope defects observed upon MSMEG_1353 depletion are therefore best interpreted as a secondary consequence of metabolic uncoupling. When central carbon metabolism and alternative carbon utilisation pathways are transcriptionally suppressed, the supply of reducing equivalents and lipid precursors becomes limiting (55). Under such conditions, continued envelope biosynthesis without appropriate regulation would be energetically unsustainable. The observed membrane destabilisation may thus reflect a failure to properly downscale or remodel envelope synthesis in response to metabolic stress, resulting in a structurally compromised cell envelope. This interpretation also provides a coherent explanation for the seemingly paradoxical antibiotic phenotypes. The emergence of isoniazid resistance despite enhanced envelope permeability is counterintuitive and therefore, suggests the involvement of additional resistance mechanisms, beyond simple changes in drug entry. However, if MSMEG_1353 depletion perturbs redox homeostasis or respiratory efficiency, the reduced activation of isoniazid becomes a plausible outcome. Additionally, the elevated mycolic acid observed in the KD strain may be responsible for the reduced susceptibility observed in the strain. In contrast, hypersensitivity to vancomycin is consistent with physical weakening of the mycomembrane, allowing access to a drug that is otherwise excluded. Consistent with this, the MSMEG_1353 KD strain exhibits a substantial reduction in PDIM, a critical determinant of mycobacterial cell envelope impermeability (56). Loss of PDIM has been shown to increase the cell wall permeability, and PDIM-deficient strains of *M. tuberculosis* display heightened susceptibility to vancomycin. Taken together, these observations support a model in which PDIM depletion underlies the enhanced vancomycin sensitivity observed upon MSMEG_1353 depletion (34, 45). The divergence between these two phenotypes reinforces the idea that MSMEG_1353 influences both metabolic physiology and envelope integrity, rather than acting solely at the level of permeability. The transcriptional and metabolomic profile of the knockdown further supports this interpretation. Rather than inducing compensatory pathways, the cells exhibit a broad metabolic contraction, including repression of glycolysis, and short-chain carbon metabolism. Such a response is characteristic of an energy deficient state in which the bacterium prioritizes survival over growth.

From an evolutionary perspective, the high conservation of MSMEG_1353 across mycobacterial species, including those with highly reduced genomes, supports that this coordination is fundamental to mycobacterial viability. Collectively, these findings position MSMEG_1353 as a previously unrecognized node linking ATP sensing, core metabolism, and cell envelope stability. This work expands the current paradigm of mycobacterial envelope biology by highlighting the importance of energy dependent regulatory proteins in maintaining envelope homeostasis. Targeting such coupling mechanisms may represent an underexplored strategy for destabilizing the mycobacterial envelope while simultaneously sensitizing cells to metabolic stress.

## 4. Materials and methods

### 4.1 Growth conditions

*E. coli* DH5α strains were employed for vector construction. Cultures were grown in Luria Bertani (LB) broth supplemented, when required, with appropriate antibiotics (kanamycin at 50 µg/ml and hygromycin at 100 µg/ml). For solid media preparation, LB broth was supplemented with 1.5% agar.

The mycobacterium strains used in this study were derivatives of *M. smegmatis* mc²155 which was cultured in Middlebrook 7H9 broth supplemented with Tween 80/tyloxapol (0.05%) and glycerol (0.25%), or on Middlebrook 7H10/7H11 agar supplemented with glycerol (0.5%). Where appropriate, antibiotics (kanamycin at 20 µg/ml and hygromycin at 50 µg/ml) and the inducer anhydrotetracycline (ATc; 100 ng/ml) were added. All cultures were incubated at 37 °C with shaking at 150 rpm.

### 4.2 Strain construction

All the cloning and initial vector constructions were done in *E. coli* following standard recombinant DNA procedures (57) before being introduced into mycobacterium. For generation of CRISPRi knockdown strains, the plasmid backbone was constructed as described earlier (24). Briefly, using a two-plasmid system CRISPRi system-PSTKiT carrying *cas12a* and pSTHT carrying guide RNA targeting MSMEG_1353 was transformed in *M. smegmatis*.

pMV261 (58) was used for cloning the MSMEG_1353-GFP over-expression strain. MSMEG_1353 was amplified from *M. smegmatis* genomic DNA with primers bearing the restriction site (BamHI and HindIII) (S.1E). Primers bearing the restriction flanks were used to amplify GFP gene from pMN437 (59) which was ligated into (ClaI and HindIII site) pMV261 to obtain pMV261-GFP. Subsequently, the gene of interest MSMEG_1353 was cloned in frame at the sites-BamHI and HindIII in pMV261-GFP to generate plasmid pMV261-1353-GFP. The construct was confirmed by Sanger sequencing and transformed into *M. smegmatis*.

### 4.3 Growth curve assay

The recombinant strains were grown to stationary phase and were used to inoculate fresh Middlebrook 7H9 medium with or without ATc at a starting O.D_600nm_ of 0.02 in a 96 well plate. The O.D_600nm_ was monitored in Tecan Pro plate reader at 37°C, every 1-2 hours for 48 hours. The O.D values were plotted against time to evaluate the fitness of strains.

For carbon sources and antibiotic treatments, compounds were dissolved in water. The pre-depleted knockdown strains were inoculated at O.D 0.02 in presence of ATc and drugs and carbon sources were used at applicable concentration (10mM). A 200 μl culture was dispensed per well in a 96-well plate in triplicates. O.D_600nm_ was evaluated using a Tecan Pro plate reader for 48 hours. A suitable time point was selected at which both strains were in the log phase for further analysis. Percent growth was calculated relative to the vehicle control for each strain for all curves, data represent the mean ± standard error for technical triplicates. Data are representative of at least two independent experiments performed.

### 4.4 ATP binding assay

ATP binding was assessed using the fluorescent nucleotide analogue 2′,3′-O-(2,4,6-trinitrophenyl) adenosine 5′-triphosphate (TNP-ATP). All samples were dissolved in Tris-HCl buffer (50 mM Tris-HCl, 50 mM KCl, pH 7.2). A working solution of TNP-ATP (0.64 mM) was prepared by diluting the 6.4 mM stock (Invitrogen) in assay buffer. Lysozyme was used as a non-ATP binding negative control and Glucokinase was used as a positive ATP-binding control, prepared at a final concentration of 4 µM. Purified protein samples were diluted in Tris-HCl buffer to a final concentration of 4 µM. In a microplate, 200 µL of buffer (blank), lysozyme control, glucokinase positive control, purified Rv0647c protein sample were dispensed into wells in triplicate, followed by addition of TNP-ATP to a final concentration of 5 µM. Plate was placed in a microplate reader, and fluorescence measurements were recorded using an excitation wavelength of 410 nm with emission scanned from 500 to 600 nm. Fluorescence data was exported for analysis, and protein specific TNP-ATP binding was calculated.

### 4.5 Electron microscopy

SEM: The knockdown strains (5 ml of 0.6 O.D cells) were harvested at 5000 *g* for 10 minutes at 4 °C and washed twice in 0.1M phosphate buffered saline, pH-8 (PBS). The cells were then fixed using 2.5% glutaraldehyde for 2 hours in dark. The fixed cells were then centrifuged and washed with PBS to remove traces of glutaraldehyde. The washed cells were subsequently dehydrated by serially treating with alcohol concentrations: 10%, 30%, 40%, 50%, 70% and 90% respectively. The cells were finally suspended in 100% alcohol and air dried to remove excess water content from the sample. The sample was drop cast on a carbon taped aluminium stub and was imaged after gold sputter coating.

SEM was performed at ×10,000 on the Zeiss SEM instrument using an accelerating voltage of 20 kV.

TEM: For TEM, a similar protocol was followed with exception of adding ruthenium red along with glutaraldehyde during fixation to increase contrast. The cells were later washed and embedded in parafilm which were sliced using microtome and visualized at 120kV. More than 20 images at different magnification and fields were documented. The representation of such image is shown in the manuscript.

### 4.6 Detection of total reactive oxygen species (ROS)

The content of total reactive oxygen species (ROS) was measured with the fluorescent probe dichlorodihydrofluorescein diacetate (DCFDA) as described earlier (60). This process was facilitated using the compound 2′,7′-dichlorofluorescein diacetate (DCFDA) dissolved in DMSO to create a main stock solution with a concentration of 20 mM, which was stored at-20°C. This stock solution was further diluted to 1 mM to prepare working stocks for analysis. For initial measurements, ATc treated or untreated mycobacteria cultures grown to the stationary phase were diluted to O.D_600_ of 0.8 in the medium and stained with a final concentration of 10 μM DCFDA. Aliquots of 100 μl were then dispensed into replicates in a dark 96-well plate with an optical flat bottom. Following a 70-minute incubation period at 37°C, fluorescence measurements were taken using a plate reader set at an excitation wavelength of 488 nm and an emission wavelength of 520 nm.

The relative fluorescence units obtained were normalised to absorbance to ensure accuracy and consistency in the data across the samples.

### 4.7 Disc assay

The knockdown strains were pre-depleted of target gene product in presence of ATc for at least 15 hours. Subsequently, the strains were harvested and washed in PBS (0.1M, pH8). The cells (200 μl at 0.D 1) were then suspended in warm 0.6% agarose and mixed well such that a homogenous suspension formed. The resuspended cells were gently poured over MB agar plates containing hygromycin, kanamycin and ATc. The discs impregnated with 5% SDS were placed gently on to the agar. The plates were incubated for 3-4 days until a visible lawn appeared and zone of inhibition was measured and tabulated.

### 4.8 Membrane polarisation assay

The knockdown and NTA were harvested and washed with PBS buffer. Subsequently, the strains were treated with and without CCCP 5 μM for 5 minutes followed by staining with DiOC2(3) dye. Stained bacterial samples were analyzed on a flow cytometer equipped with a 488 nm excitation laser. Fluorescence emission was collected in the green and red channels using filters appropriate for BB515A and PerCP-Cy5 detection, respectively. Forward scatter (FSC), side scatter (SSC), and fluorescence signals were acquired using logarithmic amplification. Membrane potential was acquired using BD flow cytometer and analysed with FlowJo software. The quantified mean fluorescence intensity of red to green fluorescence was plotted, which serves as a size independent indicator of membrane polarisation.

### 4.9 Protein localisation assay

For protein localisation studies, the cells expressing the GFP-fused protein were harvested at mid-log phase and washed twice in PBS (0.1M, pH-8). The washed cells were drop cast on a glass slide as a thin film and air dried and subsequently mounted using prolong gold antifade. The samples were subsequently imaged under 63X by confocal microscopy (MICA, Leica).

For RADA pulse chase experiment, saturated cultures of *M. smegmatis* strains bearing MSMEG_1353 GFP fused plasmid (pMV261-1353-GFP), were sub-cultured and grown to reach log phase to ensure all cells are in same phase of growth. Subsequently, cells were inoculated into fresh medium containing RADA dye at 20µM concentration at 0.02 O.D_600nm_. The cells were given a long pulse of 15 hours to ensure that all cells are uniformly stained with the dye. Subsequently, the cells were washed thrice with media and allowed to grow in absence of RADA for 3 hours during the chase period. The cells were then washed with PBS and mounted on slides using Antifade. The slides were visualised using GFP and TRITC filter (excitation: 560nm and emission: 630nm) using Leica MICA confocal microscope.

### 4.10 RNA isolation, qPCR and RNAseq

Mycobacterial cultures were harvested at an optical density of 3–5 by centrifugation at 2,000 g for 10 min. Cell pellets were resuspended in 1 ml TRIzol reagent and transferred to screw cap tubes containing lysing matrix for homogenisation. Cell disruption was performed using a bead beater (FastPrep-25 5G, MP Biomedicals) at 6.0 m s ^−1^ for 45 seconds for two cycles, with a 2 minutes ice-cooling interval between cycles. The lysates were centrifuged at 8,000 *g*, and the supernatant was transferred to fresh tubes. Total RNA was isolated using the Zymo Research RNAzol kit according to the manufacturer’s instructions.

Eluted RNA was treated with TURBO™ DNase (Thermo Fisher Scientific) for 1 hour at 37 °C to remove residual genomic DNA, followed by column purification using the Monarch DNA Cleanup Kit (New England Biolabs). Purified RNA (1 µg) was reverse transcribed using random hexamers and the Verso cDNA Synthesis Kit (Thermo Fisher Scientific) following the manufacturer’s protocol. Quantitative PCR (qPCR) was performed using SYBR Green chemistry, with cDNA as the template. Gene expression levels were normalized to *sigA* (MSMEG_2758) and relative expression was calculated using the ΔΔCt method. No-reverse transcriptase controls, no-template controls, and genomic DNA positive controls were included in each experiment. Primers were designed using the OligoAnalyzer™ Tool (IDT) and are listed in S.1E.

RNA-sequencing was outsourced to Novelgene Pvt. Ltd. Total RNA was isolated from three biological replicates of each strain. RNA quality checked samples were fragmented, and cDNA libraries were prepared according to standard protocols, followed by high-throughput sequencing. Raw FASTQ files were pre-processed using **fastp** to remove adapter sequences and low-quality reads (Phred score < 20 and read length < 50 bp). High-quality reads were mapped to the *Mycolicibacterium smegmatis* reference genome (NCBI accession CP000480.1) using **HISAT2**. Read quantification and differential gene expression analysis were performed using the DESeq2 package in R. Genes with an adjusted *P* value ≤ 0.05 and a log _2_ fold change ≥ +1 or ≤ −1 were considered significantly up-or down-regulated, respectively.

### 4.11 Metabolomics

The strains were grown to log phase and harvested after washing with ammonium buffer. Metabolites were extracted by adding five volumes of chilled 80% (v/v) methanol relative to sample weight. Samples were subjected to bead beating for 15 minutes under temperature-controlled conditions, followed by sonication for 10 minutes at 4 °C to enhance metabolite recovery. The extracts were incubated at −20 °C for 2 hours to facilitate protein precipitation and subsequently centrifuged at 13,000 *g* for 10 minutes at 4 °C. The resulting supernatant was collected and filtered through 0.22 µm filters, and 10 µl of the prepared extract was injected for analysis.

The LC–MS analysis was outsourced from Valerian chem pvt ltd using a Dionex Ultimate 3000 UHPLC system (Thermo Scientific, MA, USA) coupled to a Orbitrap Exploris mass spectrometer (Thermo Fisher Scientific, Sunnyvale, CA, USA). Data were acquired separately in positive (ESI+) and negative (ESI−) electrospray ionization modes. Chromatographic separation was carried out using solvent A (water containing 0.1% formic acid, v/v) and solvent B (acetonitrile containing 0.1% formic acid, v/v) at a flow rate of 0.300 ml/min with the following gradient: 0 min (5% B); 0–3.5 min (5–30% B); 3.5–7.5 min (30–60% B); 7.5–10 min (60–90% B); 10–12.5 min (90% B); 12.5–14 min (90–5% B); and 14–16 min (5% B). The total run time was 16 minutes.

Raw data files were processed using Compound Discoverer (version 3.2, Thermo Fisher Scientific). Compound consolidation was performed using a mass tolerance of 5 ppm and a retention time alignment window of 0.2 minute. Metabolite annotation was conducted by matching spectral features against the ChemSpider and mzCloud databases. Results were filtered based on a mass accuracy range of −5 to +5 ppm and the presence of annotated compound names. Positive and negative mode datasets were processed independently and subsequently combined, with duplicate entries removed prior to downstream analysis.

### 4.12 Total lipid extraction

MSMEG_1353 KD and NTA strains were harvested in log phase. Lipids were extracted as described previously (61). Bacterial cell pellets underwent sequential lipid extraction using chloroform/methanol mixtures in ratios of 1:2 and then 2:1 (v/v). The resulting extracts were combined, washed with water and dried. The dried lipids were then redissolved in chloroform and the spots were applied on TLC plates. Silicia gel G60 plates served as the stationary phase. For separating trehalose polyphleates (TPPs) and Glycopeptidolipids (GPLs), the mobile phase consisted of chloroform:methanol (90:10 v/v) (62). For resolving trehalose monomycolate (TMM) and Trehalose dimycolate (TDM), the mobile phase used was chloroform:methanol:water (90:10:1 v/v/v). TPP and GPLs bands were visualized by spraying the developed plates with 0.2% (w/v) anthrone in concentrated sulphuric acid, followed by charring to reveal the spots. For visualization of TMM and TDM, 10% (w/v) phosphomolybdic acid in ethanol was used followed by heating at 120°C. Band intensities were quantified using IMAGEJ software (NIH,USA), by measuring density values and normalizing against loading control.

### 4.13 Fatty acid and mycolic acid extraction

Cultures of MSMEG_1353 KD and NTA strains were grown to mid log phase (OD₆₀₀ = 0.8 to 1.5) and harvested by centrifugation at 3,500-4,000 *g* for 10 minutes at 4°C. The cell pellets were rinsed twice with sterile distilled water or PBS (pH 7.4) and then standardized to equivalent biomass (100-200 mg wet weight). Equal weight pellet were resuspended in 2 ml of tetrabutylammonium hydroxide (TBAH) and heated overnight at 100 °C to achieve full saponification and release of fatty acids and mycolic acids following cooling to room temperature, the mixture was subjected to phase transfer methylation by adding 4 ml dichloromethane, 300 µl iodomethane, and 2 ml water. The samples were vigourosly mixed for 1 hour at room temperature and then centrifuged at 3,500 *g* for 10 minutes to separate the layers. The upper aqueous phase was removed and discarded. The lower organic phase was washed thrice with 4 ml water each time, involving 15 minutes of mixing and centrifugation at 3,500 *g* for 10 minutes per wash. The washed organic phase was evaporated to dryness under gentle nitrogen stream. The dried residue was redissolved in 3 ml diethyl ether, subjected to sonication in a water bath for 10 minutes at room temperature, and centrifuged at 3,500 *g* for 10 minutes. The clear supernatant, containing fatty acid methyl esters (FAMEs) and mycolic acid methyl esters (MAMEs) was transferred to a fresh glass tube, dried under nitrogen, and finally reconstituted in 100–200 µl dichloromethane. To separate FAMEs and MAMEs, aliquots (2–3 µl) were spotted onto aluminum-backed TLC plates. Plates were developed in hexane:ethyl acetate(19:1, v/v) (63). After the solvent front reached the top, plates were air dired briefly and subjected to three successive development cycles in the same solvent system to enhance resolution of mycolic acid subclasses. Lipid bands were detected by exposure to iodine vapor and documented accordingly.

### 4.14 Apolar lipid extraction

Apolar lipids including triacylglycerols, phthiocerol dimycocerosates and other envelope components that are non-covalently associated were extracted from equal amount of biomass (200-300 mg of wet weight per sample). Cells were harvested from mid-logarithmic phase cultures (OD₆₀₀ = 0.8 to 1.5) by centrifugation at 3,500-4,000 *g* for 10 minutes at 4°C and washing them twice with sterile distilled water. Equal weight of biomass (250 mg) were suspended in 10 ml of a methanol: 0.3% (w/v) aqueous sodium chloride (10:1, v/v) in glass tubes, then 5 ml of petroleum ether (60-80 °C boiling range) was added, and the mixture was vigorously agitated at room temperature for 1 hour. The organic phases obtained from the three sequential extractions with petroleum ether were combined and subjected to centrifugation at 3,000 *g* for 15 minutes to pellet any residual insoluble material. The resulting clear supernatant was carefully decanted into a new tube and evaporated completely under gentle nitrogen stream to yield a dry lipid layer. The residue was subsequently dissolved in 100 µL of dichloromethane for further analysis. For TLC, equal volumes of resuspended extracts were spotted onto aluminium-backed silica gel 60 F₂₅₄ plates (Merck) and developed once in petroleum ether (60-80°C):diethyl ether (90:10, v/v) for PDIM (64, 65), and in hexane:diethylether:acetic acid, (70:30:1, v/v/v) for additional nonpolar lipids (i.e. TAG, DAG) (66). Plates were air dried, immersed in 10% (w/v) phosphomolybdic acid in absolute ethanol, drained of excess reagent, and heated at 120°C for 5-10 minutes to visualize lipids as blue-green spots. Band intensity was quantified densitometrically (e.g., using ImageLab software).

Apolar lipids including triacylglycerols, phthiocerol dimycocerosates, and envelope components that are non-covalently associated were extracted from equal amount of biomass. Equal weight of biomass (250 mg) were suspended in 10 ml of a methanol:0.3% (w/v) sodium chloride aqueous solution (10:1, v/v) in glass tubes, followed by addition of 5 ml of 60-80 °C petroleum ether and mixed the mixture vigorously at room temperature for at least 15 minutes. After centrifugation at 3,000 *g* for 5 min, the upper organic phase was collected, and the extraction repeated twice more (three extractions total) the pooled organic phases were centrifuged at 3000 *g* for 15 minutes to remove residual particulate matter; the clear supernatant was transferred to a clean tube, and then evaporated to dryness under a stream of nitrogen gas. The dried residue was redissolved in 100 µL of dichloromethane. For TLC analysis, equal volumes of resuspended extracts were spotted onto aluminium-backed silica gel 60 F₂₅₄ plates (Merck) and developed once in petroleum ether (60-80°C):diethyl ether (90:10, v/v) for PDIM and in hexane:diethyl ether:acetic acid, (70:30:1, v/v/v) for additional nonpolar lipids (i.e. TAG, DAG). Plates were air dried, immersed in 10% (w/v) phosphomolybdic acid in absolute ethanol, drained of excess reagent, and heated at 120°C for 5–10 minutes to visualize lipids as blue-green spots. Band intensity was quantified densitometrically.

### 4.15 Statistical analysis

All experiments were repeated two to three times with minimum of three technical replicates. Statistical analysis on various data sets were carried out using GraphPad Prism 8.0. For all graphs, mean and standard deviation are plotted. Statistical significance of the data was obtained at 95% confidence intervals for P values <0.05. Statistical significance was determined as follows: ns, not significant (p ≥ 0.05); *p < 0.05; **p < 0.01; ***p < 0.001; ****p < 0.0001.

