## SUPPLEMENTARY S.1 for "MSMEG_1353 is a critical determinant of metabolic and cell envelope homeostasis in mycobacteria"

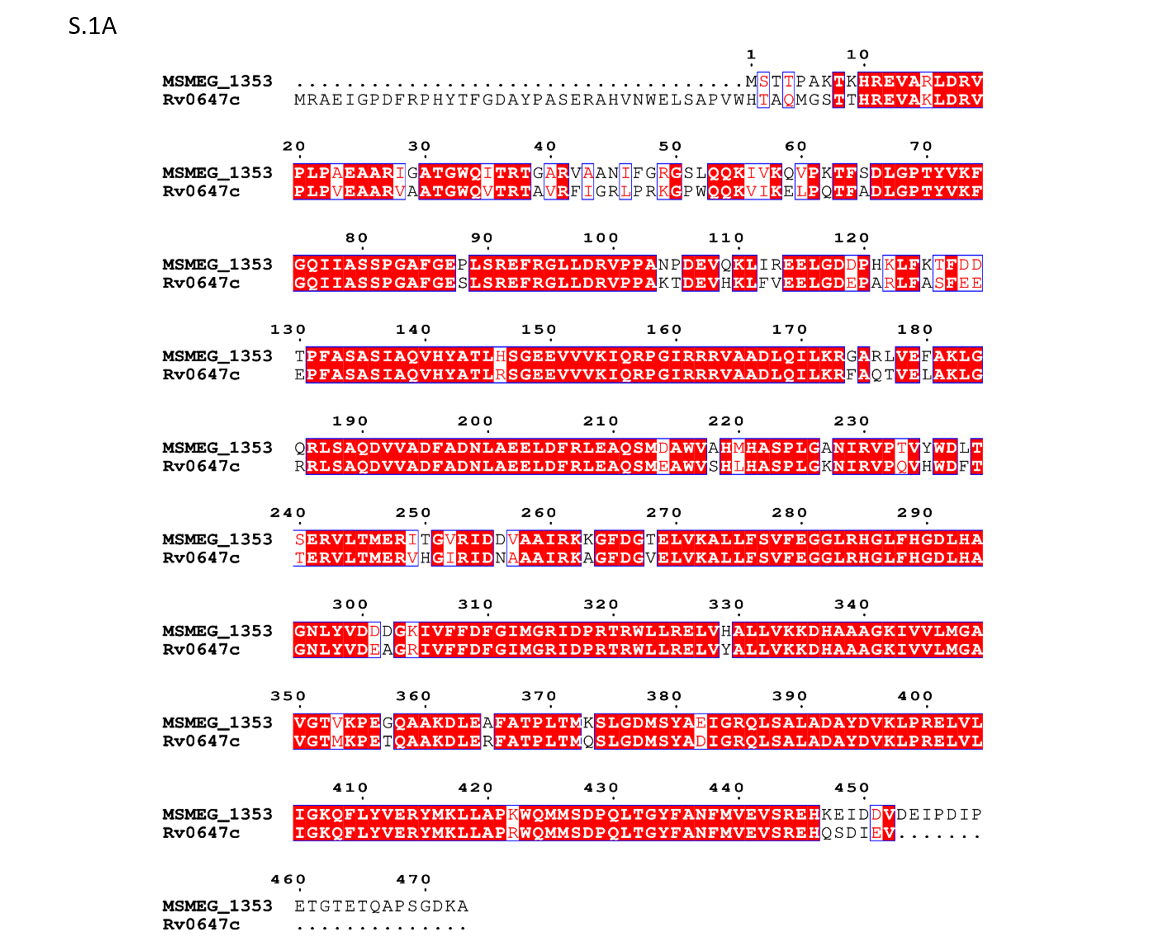

**S.1A:** Multiple sequence alignment of MSMEG_1353 and Rv0647c homologs: Alignment was generated using ClustalW and rendered in ESPript. Strictly conserved residues are boxed in red; similar residues are shown in red font using espript

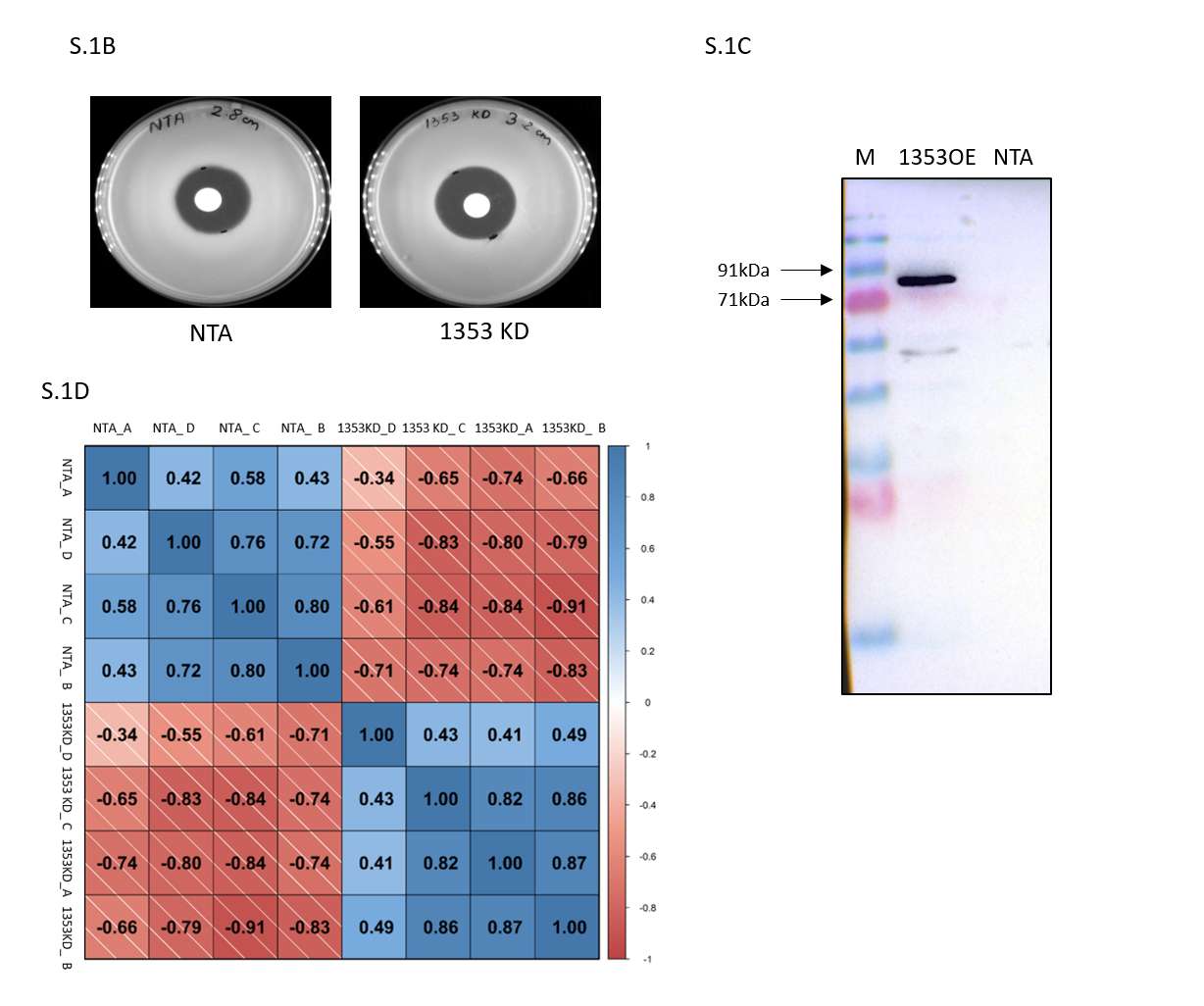

**S.1B:** Plates showing SDS sensitivity of the indicated strains upon exposure to 5% SDS measured in terms of larger zone of inhibition. **S.1C:** Western blot image showing the migration of 1353 GFP fusion protein at the desired size **S.1D:** Metabolomics correlation plot of MSMEG_1353 KD and NTA.

| **Oligonucleotide description** | **Primer Name** | **Sequence (5'-3')** |
| --- | --- | --- |
| Primers used for gene expression  check by qPCR | MSMEG_1393 RTF | TCTGACACCAACACGCCTTC |
|  | MSMEG_1393 RTR | CTT GCT GAC CGA GTT GCC |
| Primers used for gene expression  check by qPCR | MSMEG_6629 RTF | GCT CGC ATC GAC ACG AAA CG |
|  | MSMEG_6629 RTR | CGTTGATGTTGGCGATCCACAC |
| Primers used for gene expression  check by qPCR | MSMEG_4920 RTF | GGTTCGCTCAAGGATTTCTCC |
|  | MSMEG_4920 RTR | CGA CAG GCA CAT CTT GTT GAT |
| Primers used for gene expression  check by qPCR | MSMEG_6744 RTF | TATGCCAACTGCCCCAAGTACA |
|  | MSMEG_6744 RTF | CCC AGT CGA CGA ACA GCA AC |
| Primers used for gene expression  check by qPCR | MSMEG_3199 RTF | ACACCATCGTGTTCTGCGG |
|  | MSMEG_3199 RTR | AGT TCG TCG CCG TTG ATG C |
| Primers used for gene expression  check by qPCR | MSMEG_0520 RTF | CGTTGCTCGCAGGCACCAC |
|  | MSMEG_0520 RTR | TGC CGT GCG CGT TGG ACA C |
| Primers used for reference gene expression  check by qPCR | MYS GENOTYP RTF | CCGAAGAGCTCGCCAAGGAG |
|  | MYS GENOTYP RTR | GTCACCGAGCTGGCTGTCAC |
| Primers used for gene expression  check by qPCR | 1353 RTF | CAACATCTTCGGCCGCG |
|  | 1353 RTR | GCA CCT CGT CCG GGT TC |
| Oligos used for cloning the guide RNA targeting MSMEG_1353 | MSM 1353 TOP GRNA | TAGAT GGCCGCGGATCTCTTCAGCAGAA |
|  | MSM 1353 BOTTOM GRNA | AGAC TTCTGCTGAAGAGATCCGCGGCCA |
| Oligos used for cloning the Non targeting guide RNA | NTA TOP | TAGATACCAAGGACACATTCGAGCTCT |
|  | NTA BOTTOM | AGACAGAGCTCGAATGTGTCCTTGGTA |
| Primers used for cloning of MSMEG_1353 in GFP fusion overexpression strain | GFP CLA R | GTCGTCCTTGTAGTCCTTGTACAGCTCGTCCATGC |
|  | GFP HIND F | ATGTAAGCTTATGTCGAAGGGCGAGGAGCT |
|  | MSMEG_1353 R | TAT T AAGCTT GG CCT TGT CCC CGCTG |
|  | 1353 BAMH +1 F | ATT AGG ATC C A AT GAG CAC TAC GCC GG |

**S.1E:** List of oligos used in the study
